# Predicting drug polypharmacology from cell morphology readouts using variational autoencoder latent space arithmetic

**DOI:** 10.1101/2021.09.02.458673

**Authors:** Yuen Ler Chow, Shantanu Singh, Anne E. Carpenter, Gregory P. Way

**Author notes:** **Corresponding author:** G.P.W.

## Abstract

A variational autoencoder (VAE) is a machine learning algorithm, useful for generating a compressed and interpretable latent space. These representations have been generated from various biomedical data types and can be used to produce realistic-looking simulated data. However, standard vanilla VAEs suffer from entangled and uninformative latent spaces, which can be mitigated using other types of VAEs such as β-VAE and MMD-VAE. In this project, we evaluated the ability of VAEs to learn cell morphology characteristics derived from cell images. We trained and evaluated these three VAE variants—Vanilla VAE, β-VAE, and MMD-VAE—on cell morphology readouts and explored the generative capacity of each model to predict compound polypharmacology (the interactions of a drug with more than one target) using an approach called latent space arithmetic (LSA). To test the generalizability of the strategy, we also trained these VAEs using gene expression data of the same compound perturbations and found that gene expression provides complementary information. We found that the β-VAE and MMD-VAE disentangle morphology signals and reveal a more interpretable latent space. We reliably simulated morphology and gene expression readouts from certain compounds thereby predicting cell states perturbed with compounds of known polypharmacology. Inferring cell state for specific drug mechanisms could aid researchers in developing and identifying targeted therapeutics and categorizing off-target effects in the future.

## 1. Introduction

A variational autoencoder (VAE) is a generative model that can generate realistic simulated data (Kingma & Welling, 2013). As an unsupervised model, a VAE is data-driven and learns by reconstructing input data rather than by minimizing classification error as in a traditional supervised neural network. A VAE compresses data into a lower-dimensional representation and then decodes it back into the original dimensions. The compressed lower dimensional space is often referred to as a “latent space”.

The so-called “vanilla” VAEs i.e. VAEs as they were originally formulated (Kingma & Welling, 2013), are the standard choice of variational autoencoders, and have provided an important foundation in the use of generative deep learning models. However, researchers have recently identified areas for improvement, and have thus modified VAE’s loss function to overcome these issues. For example, modifying the contribution of the KL divergence term encourages disentangled latent space features, as in β-VAE (Higgins *et al*, 2016). Another variant, the so-called InfoVAE or MMD-VAE, replaces the KL divergence term with maximum mean discrepancy (MMD) to improve the model’s ability to store more information in the latent space (Zhao *et al*, 2019).

Recently, VAEs have been successfully trained on various biomedical data modalities such as bulk and single-cell gene expression data (Xue *et al*, 2020) from different assays measuring cell line perturbations and patient-derived tissue (Lotfollahi *et al*, 2019; Rampášek *et al*, 2019; Lopez *et al*, 2018; Way & Greene, 2018), DNA methylation (Levy *et al*, 2020), and cell image pixels (Lafarge *et al*, 2018; Ternes *et al*, 2021). β-VAEs have been used to produce disentangled latent representations of single cell RNA-seq data (Kimmel, 2020). Similarly, MMD-VAEs have helped retain information in the latent space in single-cell data analysis of mass cytometry and RNAseq (Zhang, 2019).

One powerful application of VAEs and other generative models is the ability to simulate meaningful new samples using an approach called latent space arithmetic (LSA). A relatively simple approach, LSA uses a series of additions and subtractions of specific sample groups in their mean latent space representations to generate synthetic samples containing specific patterns captured via arithmetic. For example, LSA has been performed using a Deep Convolutional Generative Adversarial Network (DCGAN) model to generate new representations of faces, generating realistic but synthetic images in an LSA experiment: images of men with glasses - images of men without glasses + images of women without glasses = images of women with glasses (Radford *et al*, 2015). Similarly, LSA using a generative model trained on fluorescent microscopy images, CytoGAN, predicted how cell images would look with increased nucleus size and increasing amounts of β-tubulin (Goldsborough *et al*, 2017).

Because of the success of VAEs on these various datasets, we sought to determine if VAEs could also be trained using cell morphology readouts (rather than directly on images), and further, to carry out arithmetic to predict novel treatment outcomes. To see how VAE modeling ability compared across different data types, we also trained and evaluated VAEs on two other datasets: 1) the same cell morphology data, but using all replicates instead of collapsed perturbation-specific signatures, and 2) gene expression data of the same perturbations. For our two cell morphology datasets, we used publicly-available cell morphology readouts from a Cell Painting experiment of 10,368 drug profiles from the Drug Repurposing Hub (Corsello *et al*, 2017). Our gene expression data comes from the same perturbations as measured by the L1000 assay, which quantifies mRNA transcript abundance of 978 landmark genes (Subramanian *et al*, 2017). We trained a Vanilla VAE, β-VAE, MMD-VAE, and Principal Components Analysis (PCA) on each of these datasets to compare their performance in terms of reconstruction, latent space interpretability, and predicting polypharmacology representations from these datasets (**Figure 1**). Understanding the behavior of VAE architectures in modeling morphology representations will help us to interpret various signals of morphology systems biology. Characterizing drug MOAs is a crucial step to understanding how drugs work, and for monitoring and developing new therapeutics (Chandrasekaran *et al*, 2021).

**Figure 1.**
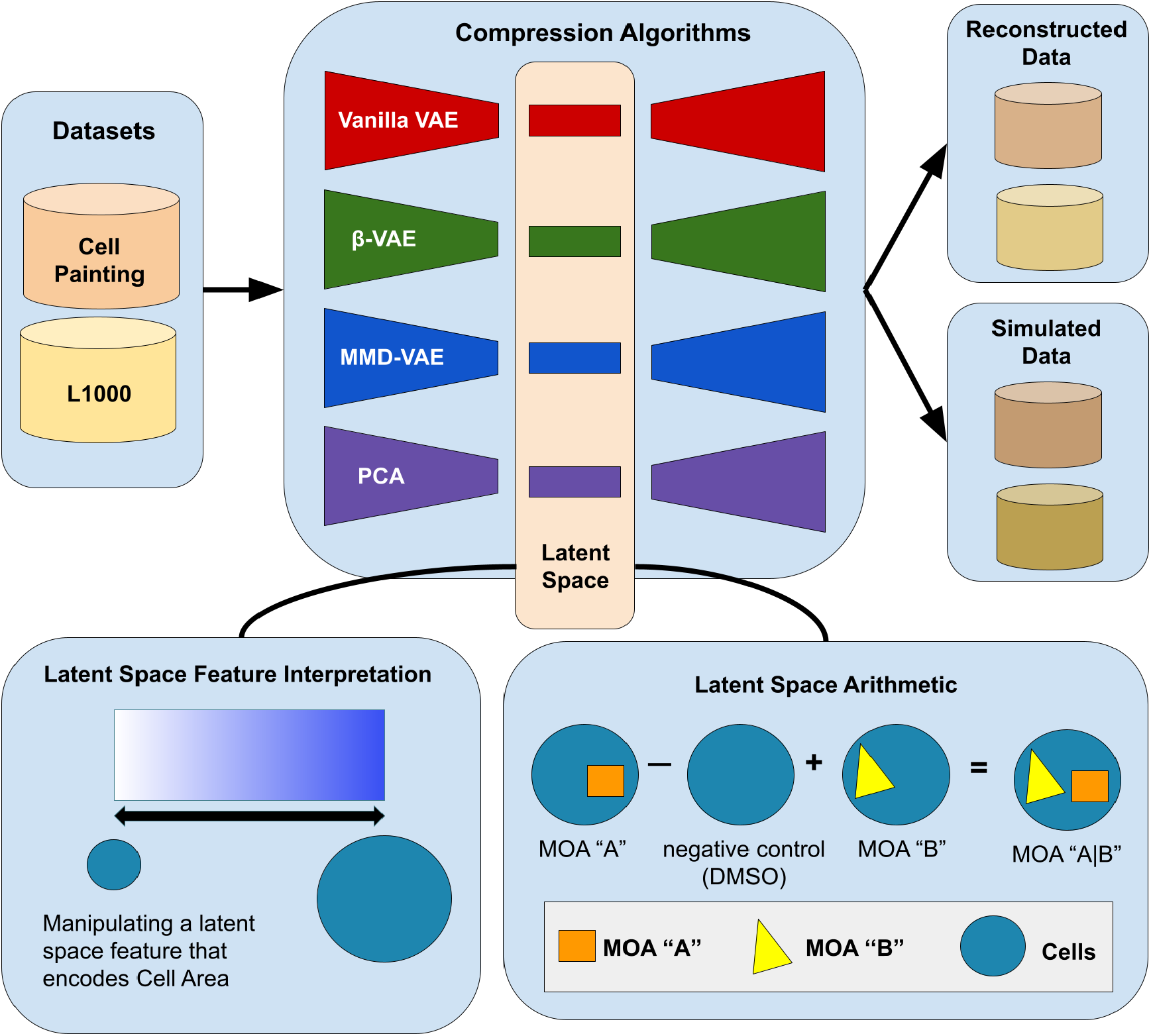
Our variational autoencoder (VAE) implementation framework, applied to determining the phenotype of cells. *One application is to predict the phenotype of cells treated with compounds that have two mechanisms of actions (MOA), given the phenotype of cells treated with compounds that have each of those single MOAs (bottom right). A VAE encodes input data into a lower-dimensional latent space and then decodes the representation back into the original data dimensions. Our data contained measurements for 588 morphology features, each averaged for each population of cells treated with a given chemical compound. Following a sweep to select optimal hyperparameters (see Methods), we set our latent space dimension to 10 dimensions. The vanilla VAE learns by minimizing a reconstruction and KL-divergence loss. The other VAE variants we tested minimize loss functions that encourage disentangled features that promote interpretability and data simulation.*

## 2. Results

### 2.1 Training variational autoencoders on cell morphology readouts

We trained our unsupervised learning models using data representing morphology readouts of A549 lung cancer cells treated with 1,571 compound perturbations from the Drug Repurposing Hub across 6 doses. Specifically, we used processed Cell Painting consensus signatures (level 5 data) from the Library of Integrated Network-Based Cellular Signatures (LINCS) (Natoli *et al*, 2021). Many of these drugs have annotations for their molecular targets and mechanisms of action (MOA). We split our input data into 80% training, 10% validation, and 10% test data balanced by plate and performed a Bayesian hyperparameter optimization using hyperopt (Bergstra *et al*, 2013).

Using optimal hyperparameters, we trained three types of VAEs: Vanilla VAE, β-VAE, and MMD-VAE and observed lower loss across epochs in real data compared to randomly permuted data (by independently shuffling the rows of each column, thus removing all correlation between features). This indicates that our VAEs have learned the data distribution by understanding the correlation between features because performance is worse when we remove correlation structure (**Figure 2**). We observed similar trends with the Cell Painting replicate profiles (level 4 data) and the L1000 datasets **(Supplementary Figure 1, Supplementary Figure 2)**. We provide latent space embeddings for all profiles in our github repository https://github.com/broadinstitute/cell-painting-vae (Chow & Way, 2021).

**Figure 2.**
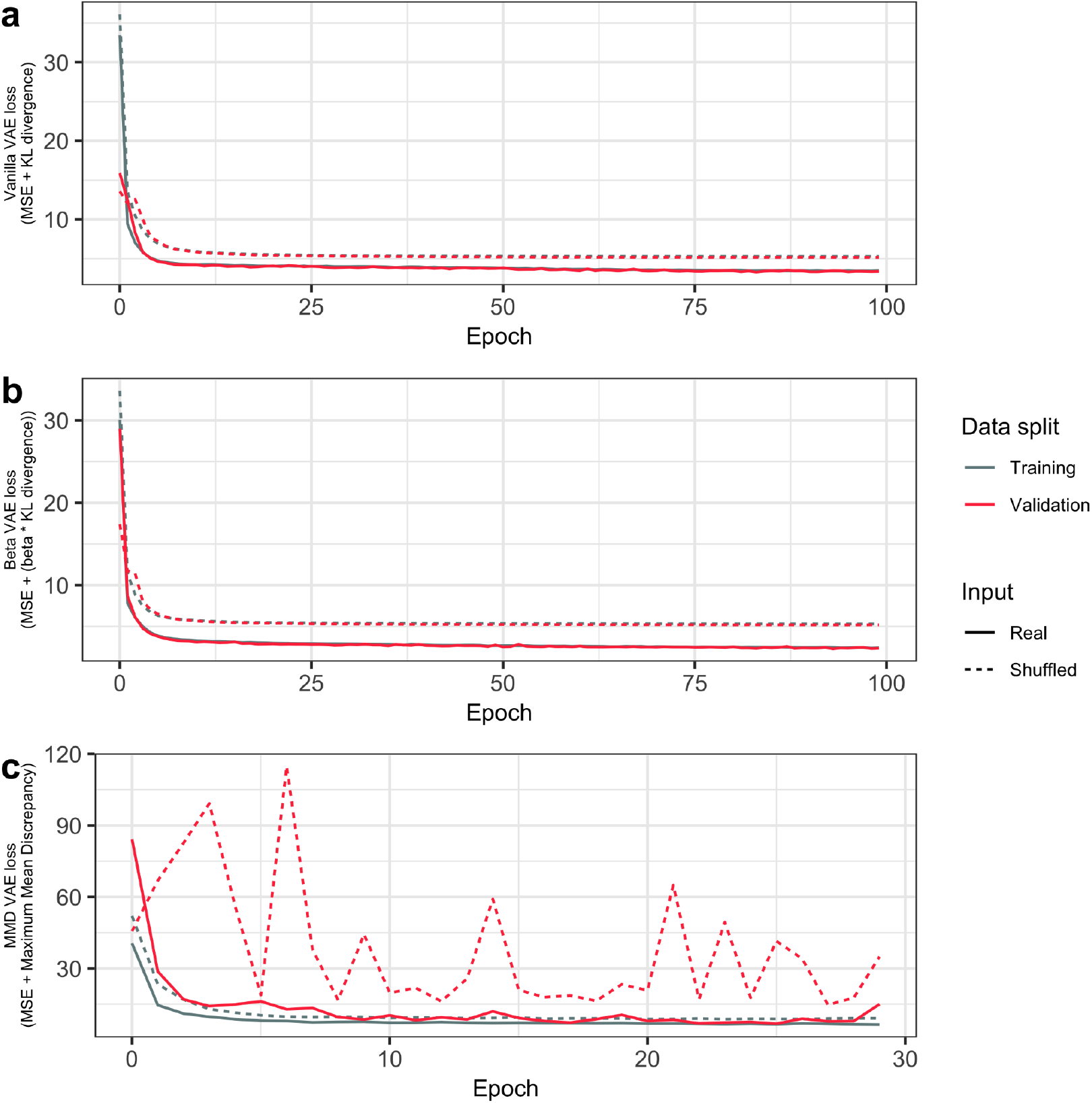
*Training curves for our VAE for Cell Painting level 5 data demonstrate a decreasing loss function for all VAE variants in Cell Painting consensus signatures from Drug Repurposing Hub compound perturbations*. We show training and validation curves for real and shuffled data in three VAE variants: **(a)** Vanilla VAE, **(b)** β-VAE, and **(c)** MMD-VAE.

Each VAE variant learns by minimizing different loss functions, which provide different constraints and learn different latent space representations (see Methods for more details). When training the β-VAE, we observed that too high a β resulted in an uninformative latent space. The decoder did not fully utilize the latent code to reconstruct input samples. On the other hand, too low a β resulted in an entangled latent space, reducing the ability to interpret latent space features and perform the LSA experiment. So, we determined the optimal β by using a method that reduces the similarity between simulated and real data (see Methods). The ability to simulate data points requires a balance of reconstruction and disentanglement, hence finding the optimal value of β that results in the best simulation would likely improve performance in the LSA experiment.

Next, we analyzed the ability of our trained VAEs to reconstruct individual samples and to simulate data. In two-dimensional Uniform Manifold Approximation and Projection (UMAP) (McInnes *et al*, 2018) embeddings, we observed that real data overlapped with both reconstructed and simulated data indicating the ability of our models to reliably approximate the underlying morphology data generating function **(Figure 3)**. Both reconstructed and simulated data did not span the full original data distribution, but were more constricted in the Vanilla-VAE compared to β and MMD-VAEs. By eye, β-VAE clearly performed the best. Our VAEs were also able to similarly reconstruct and simulate Cell Painting level 4 and L1000 data **(Supplementary Figure 3, Supplementary Figure 4)**. We also observed a tradeoff between the VAEs ability to reconstruct samples and disentangle features, as indicated by improved reconstruction but higher latent space feature correlations in β-VAE compared to Vanilla VAE **(Supplementary Figure 5)**.

**Figure 3.**
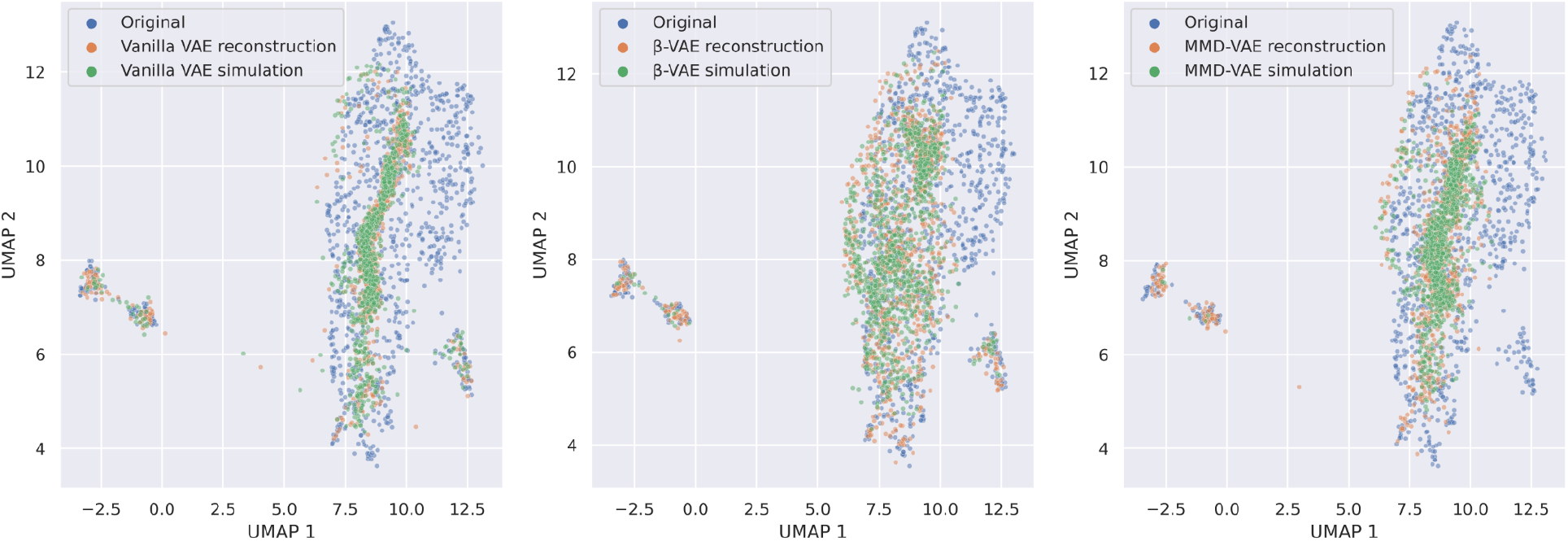
*Two-dimensional UMAP embeddings of original, reconstruction, and simulated data for Cell Painting level 5 consensus signatures in the test set*. We fit UMAP using only the original test set data and transformed the reconstructed and simulated data into this space. We simulated data by sampling from a unit Gaussian with the same dimensions as the latent space, using the same number of points as samples in the test set.

### 2.2 Interpreting Cell Painting latent space feature representations

As part of training, VAEs use different combinations of input features to generate representations. Specifically in our data, these so-called “representations” are, in essence, different combinations of morphology features that best capture signals from the input Cell Painting data. To facilitate interpretation and to understand the contribution of all morphology features to each latent space feature, we performed the following procedure: 1) Simulate +3 standard deviations of activity from one latent space feature while fixing all other latent features to zero, 2) Simulate -3 standard deviations of activity from the same latent space feature while fixing all other features to zero, 3) Pass both of these extreme-in-one-latent-feature latent spaces through the trained decoders, and 4) Subtract these two reconstructions from each other. Effectively, this procedure systematically implicates the most influential morphology features per latent space feature. This approach is similar to investigating specific VAE weight matrices (similar to PCA “loadings”) but it does not require us to set a threshold defining significant morphology feature contributions per latent feature.

As expected, we observed that our β-VAE learned more active latent features across a wide variety of morphology feature groups, compared to our Vanilla VAE and a baseline PCA. We also noticed that unlike the β-VAE where 5 of the 10 features encoded little information, all columns in our MMD-VAE were active, which indicates a more informative latent space that uses a wider diversity of morphology feature categories (**Supplementary Figure 6**).

Focusing on the MMD-VAE features, we observed that many individual latent space features encoded specific image channels and cell compartments (**Figure 4**). For example, latent feature 0 most strongly encoded Nuclei-Mito features (morphology features derived from the nucleus, specifically from the mitochondria fluorescent marker), feature 1 most strongly encoded Cytoplasm-DNA, and feature 2 most strongly encoded Nuclei-AGP and Nuclei-DNA features. The ability of the MMD-VAE to isolate these specific signals in an unbiased manner provides evidence that each image channel and compartment encodes unique information, and that these features can be used to interpret perturbation mechanisms in the future.

**Figure 4.**
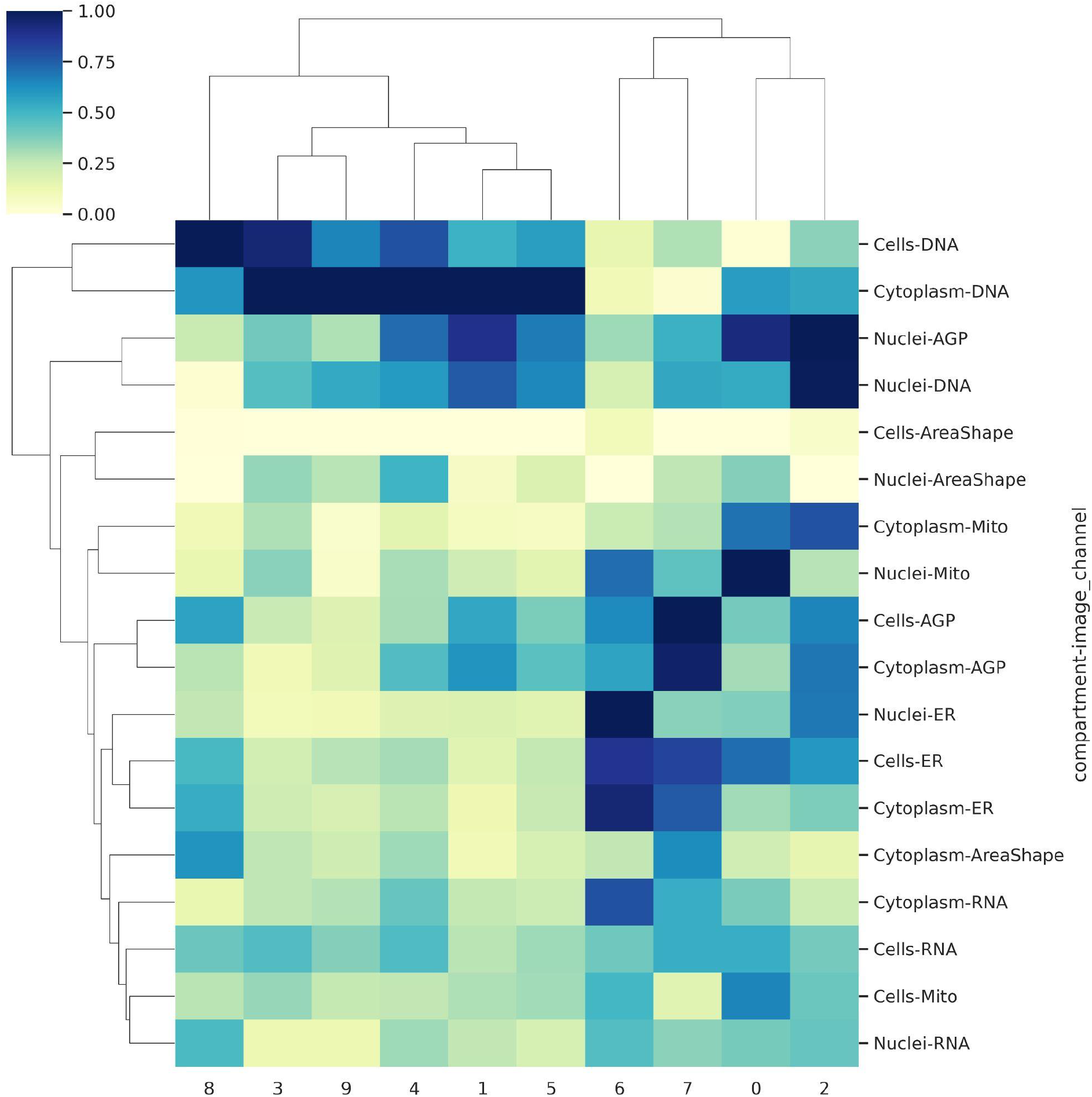
*Investigating the contribution of CellProfiler feature groups (by compartments and image channels) on individual MMD-VAE latent space features*. The dendrogram represents a hierarchical clustering algorithm applied to both rows and columns. Each color represents the mean contribution of each CellProfiler feature group to the given latent space feature normalized by column (see Methods for complete details).

### 2.3 Predicting polypharmacological cell states with latent space arithmetic

The Drug Repurposing Hub has annotated the mechanism of action (MOA) of almost all compound perturbation profiles in the LINCS Cell Painting dataset (Corsello *et al*, 2017). The MOAs represent experimentally-derived classifications indicating the most likely biological mechanism(s) of the compound. Many compounds were annotated with a single MOA, but about 14% of these compounds (214 / 1570) were annotated with two MOAs (denoted using the form “A|B”), indicating known polypharmacology; that there is evidence the compound acts through at least two separate mechanistic pathways, a common property even for marketed drugs (Proschak *et al*, 2019; Chandrasekaran *et al*, 2021). In all, the Cell Painting dataset included 84 different MOAs.

We hypothesized that we could predict cell morphology readouts for samples with MOA of the form “A|B” by performing so-called latent space arithmetic (LSA). The approach uses mean profiles of all compounds annotated with MOAs “A”, and “B”, as well as the negative control DMSO (“D”). DMSO is the solvent the experimentalists use to dissolve all drugs and therefore serves as a good profile baseline.

We predicted that subtracting the mean DMSO from the mean MOA “A” in the latent space (“A” - “D”) would allow us to obtain the essential latent space information for a profile labeled with MOA “A”. Then, adding the mean representation of latent space values for profiles with MOA “B” would allow us to obtain a compressed representation for polypharmacology profiles labeled with MOA “A|B”. We could then pass this latent representation through a VAE decoder to obtain a predicted cell with an “A|B” cell state. Taken together, our latent space arithmetic equation hypothesis is “A” - “D” + “B” = “A|B” **(Figure 5)**. We performed these analyses using all the data, including training and test sets because our test set did not contain enough variety in samples to perform LSA.

**Figure 5.**
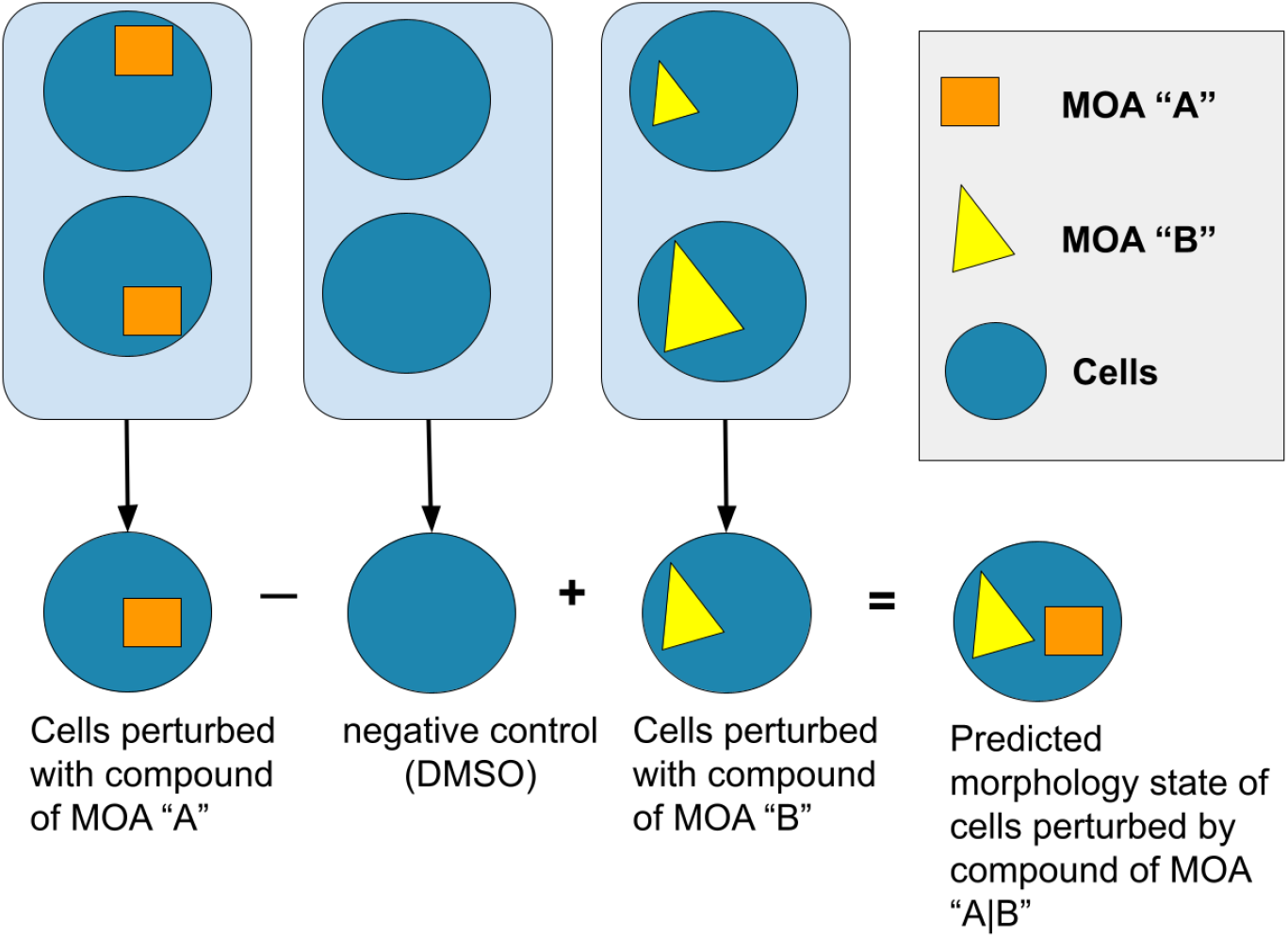
A cartoon example of the latent space arithmetic (LSA) approach we performed using consensus profiles with two annotated mechanisms of action (MOA) in the LINCS datasets.

To evaluate LSA performance, we calculated the L2 distances between the predicted and the actual “A|B”. The mean L2 distance for VAEs with unshuffled MOAs was lower than shuffled data, lower than the original input dimensions, and lower than PCA, indicating that, on average, VAEs were better at predicting polypharmacological cell states **(Figure 6, Supplementary Figure 7)**. Of the different VAE architectures, MMD-VAE performed the best in all datasets. Importantly, we also observed improved predictability for MOAs that had a greater distance away from the mean Cell Painting feature values, indicating that MOAs are easier to predict when they have a more unique phenotype, further providing support for our ability to reveal true polypharmacology cell states **(Supplementary Figure 8)**.

**Figure 6.**
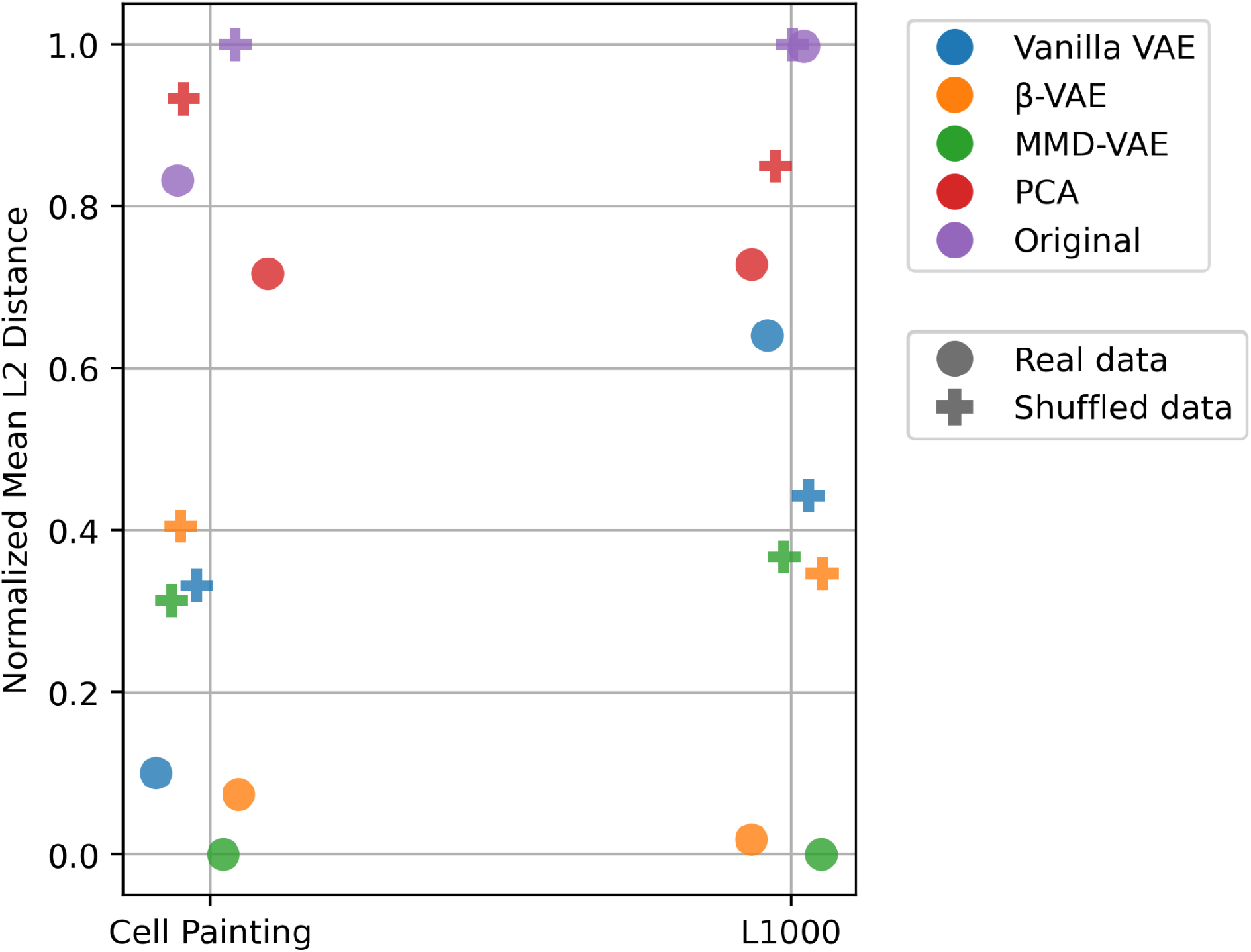
Mean *L2 distance (lower is better) between real and predicted profiles annotated with known polypharmacology (“A*|*B”) mechanisms of action (MOAs) for three different VAE architectures, PCA, and original input space*. We show results for real and shuffled data across the two LINCS datasets. To enable a more meaningful and interpretable view, we zero-one normalized the L2 distances for each dataset. Each dot represents the mean L2 distance (values are normalized within each dataset) when LSA is performed using a specific model on a specific dataset.

### 2.4 Evaluating specific polypharmacology MOA predictions

We annotated which specific MOA combinations performed best with LSA. To do so, for each polypharmacology “A|B”, we calculated a standard score (calculating the number of standard deviations from the mean), which compares 1) the L2 distance between the real and the predicted “A|B” value against 2) a randomly sampled distribution of the L2 distances between the real and predicted “A|B” values with (see methods). MOA combinations that yielded negative test scores and low p-values indicated that they were predicted better than random. We predicted polypharmacology morphology states better than the shuffled distribution for the majority of MOAs. Repeating this procedure with L1000 gene expression profiles, we observed that gene expression and morphology assays predicted many of the same polypharmacology MOAs, albeit with several exceptions **(Figure 7)**, consistent with recent work (Natoli *et al*, 2020).

**Figure 7.**
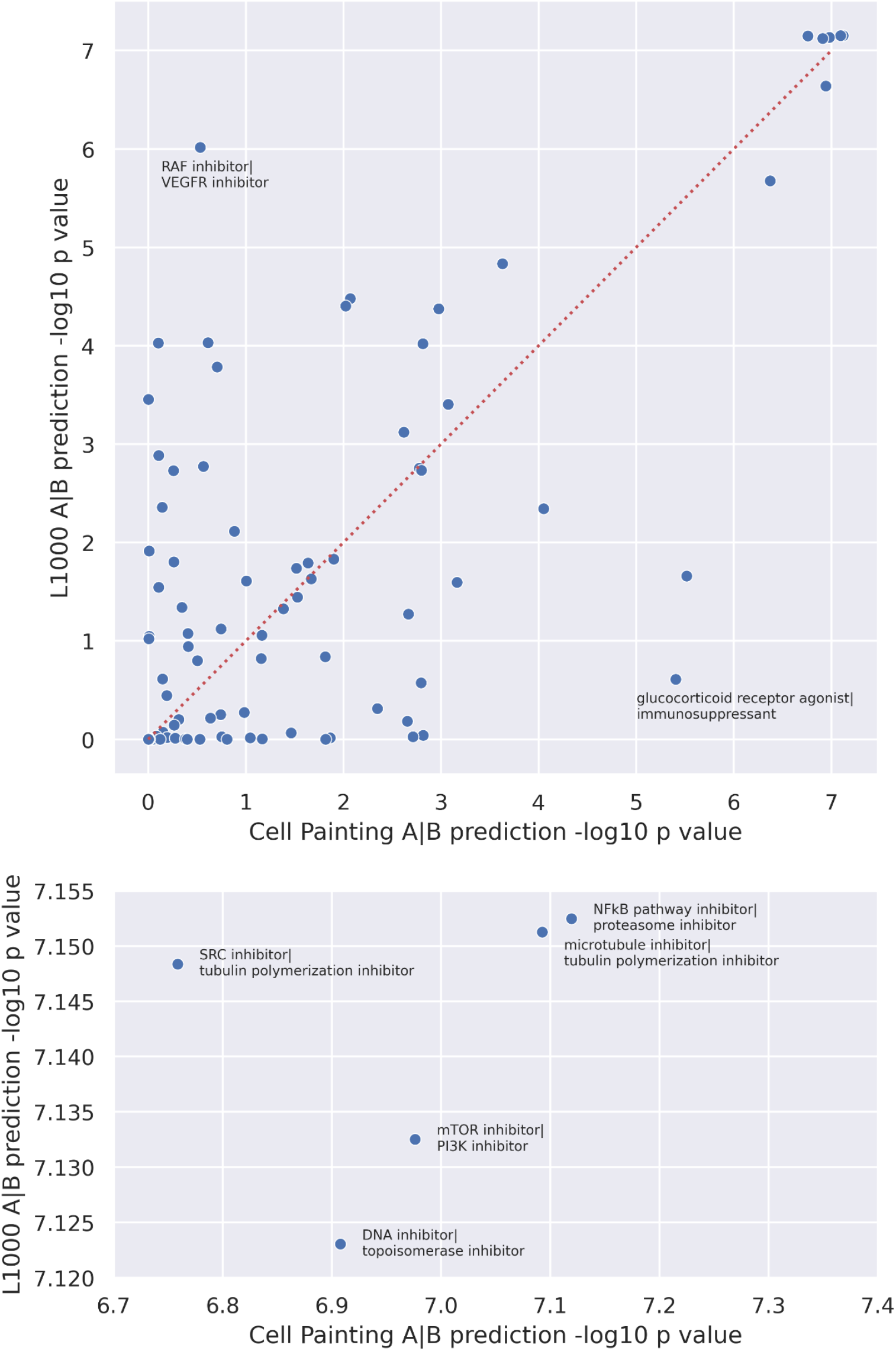
*Scatterplot of L1000 vs Cell Painting MOA performance for β-VAE, with outliers (>3 stds) excluded*. Performance is determined by the test scores between the L2 distance between predicted and real profile and the distribution of L2 distances from shuffling MOAs 10 times. *Top* - all “A|B” combinations; *bottom* - only the top 5 “A|B” with labels

One limitation of our approach was that samples in our training data were annotated with many of the same MOAs as samples used in the LSA experiment, which could potentially leak information into our evaluation set and artificially inflate performance. For our work with predicting MOAs to have practical application, we needed to know that we can predict MOAs that have never been seen by the training set. So, we removed compounds annotated to the top five best-predicted MOAs for each LSA evaluation and retrained VAEs for all VAE types. We observed that while sometimes the MOAs exhibited decreased performance, in general, those five MOAs still remain among the best predicted out of all MOAs (**Supplementary Figure 9**). This reveals two things. First, it shows that the VAEs are stable; the same MOAs are still well-predicted when we retrain the model. Second, it shows that the VAEs do not need to have to have seen the specific cell states perturbed with compounds annotated with specific MOAs to predict their cell state and we are indeed not overfitting to the data.

## 3. Discussion

Cell morphology provides an underexplored systems biology perspective of diseased and perturbed cell states. One current bottleneck in expanding this research avenue is the lack of generalizable, expressive, widely available, and interpretable morphology feature representations. Here, we used VAEs to model cell morphology representations of thousands of perturbed cell states. We determined that VAEs can be trained on cell morphology readouts rather than directly using the cell images from which they were derived. This decision comes with various trade-offs. Compared to cell images, cell morphology readouts as extracted by image analysis tools (e.g. CellProfiler) are a more manageable data type; the data are smaller, easier to distribute, substantially less expensive to analyze and store, and faster to train (McQuin *et al*, 2018). However, it is likely some biological information is lost, because these tools might fail to measure all morphology signals. The so-called image-based profiling pipeline also loses information, by nature of aggregating inherently single-cell data to bulk consensus signatures (Caicedo *et al*, 2017). Nevertheless, we successfully modeled cell morphology readouts from CellProfiler-derived representations of Cell Painting data using VAEs, and we demonstrated the power of these representations by simulating realistic-looking data, by deriving meaningful and disentangled morphology representations, and by predicting polypharmacology in certain compounds.

Using cell morphology readouts, we trained three different VAE architectures, each with different strengths and weaknesses. As expected, we observed improved modeling ability, as indicated by a lower reconstruction loss and improved data simulation, for a β-VAE model compared to a Vanilla VAE. Similarly, by using MMD-VAEs, which replaces the KL divergence term with one that calculates divergence for the full distribution of all latent features instead of each individual feature, we achieved better reconstruction than the vanilla VAE. However, the MMD-VAE failed to simulate all cell morphology modes, which the other VAE variants successfully captured. By training these VAE variants on L1000 gene expression readouts resulting from the same set of compound perturbations, we observed large differences in optimal hyperparameters as compared to training on Cell Painting image-based data (Subramanian *et al*, 2017; Xue *et al*, 2020). These observations indicate that the KL divergence penalty strongly influences cell morphology modeling ability, and that lessons learned by modeling other biomedical data types, such as gene expression, will not necessarily directly translate to cell morphology (Yang *et al*, 2021).

Through a deep inspection of the latent space features, we observed that the MMD-VAE learned the most informative representation (compared to vanilla and β-VAE), and, interestingly, uses information from all categories of Cell Painting readouts. The Cell Painting assay uses six different fluorescent stains to mark eight organelles: nuclei, endoplasmic reticulum, nucleoli, cytoplasmic RNA, actin, Golgi, plasma membrane, and mitochondria (Bray *et al*, 2016). Subsequently, image analysts use software to segment cells to distinguish nuclei from cytoplasm and measure a broad set of hand-engineered, classical image features for each cell compartment across all five fluorescent channels (McQuin *et al*, 2018). While other emerging approaches for segmenting cells and extracting morphology features exist, some based on deep learning (Lucas *et al*, 2021), the hand-engineered classical features in the profiles we used remain much more common, and are currently the most directly interpretable. It remains uncertain if all Cell Painting fluorescent stains encode important, non-redundant information or if instead some could be acquired in simpler microscopy assays (Ounkomol *et al*, 2018). Here, we provide evidence that the different stains do encode independent biological signals, by nature of the unsupervised MMD-VAE disentangling these groups. However, our view of the biological signals embedded in the latent space representations is still limited; latent space features are likely encoding morphology signatures that extend beyond the individual feature categories we quantified (e.g. Cells-AGP), and represent higher order biological processes that are interacting between cell compartments and across image channels. Nevertheless, our optimal latent space dimension included only ten features, which is typically far fewer than other modalities such as gene expression (Zhou & Altman, 2018; Way *et al*, 2020; Natoli *et al*, 2021).

Polypharmacology occurs when drugs interact with multiple targets, and it is a challenging aspect of drug discovery important for designing more effective and less toxic compounds (Reddy & Zhang, 2013). By predicting cell states of polypharmacology compounds, we can infer toxicity and simulate the mechanisms of how two compounds might interact. Using established properties of generative models (Radford *et al*, 2015), we tested our three VAE variants in a so-called latent space arithmetic (LSA) experiment to predict polypharmacology cell states. Our results indicated that LSA worked best for the MMD-VAE architecture as compared to PCA and negative control baselines. MMD-VAEs allow for both disentanglement and a meaningful latent code, which supports LSA performance. While we observed only a slightly better LSA performance for all MOAs using randomly shuffled models compared to real data, there are several possible explanations. First, MOA annotations are often noisy, unreliable, and they change over time as scientists generate new knowledge about compounds (Cox *et al*, 2020). It is also possible that Cell Painting readouts may not capture certain MOAs that specifically manifest in other modalities. Indeed, we observed that certain polypharmacology target combinations were better predicted using gene expression readouts. Finally, we observed that LSA only performs well on certain MOA combinations, with most MOA combinations showing a negligible difference from its shuffled control. This would dampen significance in total when comparing performance between shuffled and real data for certain MOAs.

## 4. Conclusion

Cells are the building blocks of life, and they change when exposed to perturbations. There are many ways to measure, describe, and interpret how these cells change. In our analysis, we found that morphological cell states, as derived from microscopy images, can be modeled through VAE unsupervised learning to reveal biological insights. We found that several of our VAE models could reconstruct and simulate morphology data with high fidelity to ground truth perturbed cell states. When analyzing the latent code, we found that each latent space feature encoded different combinations of Cell Painting features. These feature representations were unique across different VAEs, with the MMD-VAE encoding the most information across Cell Painting channels and cell compartment feature groups. Our VAE models were able to simulate not only morphology but also gene expression cell states of compounds with multiple targets. Specifically, we simulated these polypharmacology cell states better than randomly shuffled and PCA controls. Several polypharmacology cell states performed better than others, with different performance for different MOA combinations for gene expression and morphology measurements. In the future, we could use unsupervised learning and mechanism predictions to interpret cell state mechanisms in different biological modalities, predict unknown MOAs, and characterize potential off-target effects in drug discovery and treatment. We provide all software, data, and results in an open source github repository located at https://github.com/broadinstitute/cell-painting-vae (Chow & Way, 2021).

## 5. Methods

### 5.1 Data acquisition

Previously, the Connectivity Map team at the Broad Institute of MIT and Harvard perturbed A549 cells with 1,515 different drug perturbations across six different doses as part of the Library of Integrated Network-Based Cellular Signatures (LINCS) consortium. They measured cell responses to these perturbations using the L1000 and Cell Painting profiling assays. We downloaded the publicly available LINCS Cell Painting dataset (Natoli *et al*, 2020) and the publicly available LINCS L1000 data (Subramanian *et al*, 2017).

Briefly, Cell Painting is a fluorescent microscopy assay that uses a set of six unbiased dyes to mark DNA content, nucleoli, cytoplasmic RNA, endoplasmic reticulum (ER), actin, Golgi, plasma membrane, and mitochondria (Gustafsdottir *et al*, 2013; Bray *et al*, 2016). Briefly, L1000 is a bead-based gene expression assay that measures mRNA expression (Subramanian *et al*, 2017).

### 5.2 Data processing

For the Cell Painting assay, we previously applied an image analysis pipeline using CellProfiler. Previously, we used CellProfiler to segment and measure morphology features from single cells (McQuin et al. 2018). We then applied an image-based profiling pipeline, in which we aggregated and normalized single cells into compound profiles by dose per replicate (Caicedo *et al*, 2017). We performed feature selection narrowing the initial 1,789 features to 584. We used four criteria for feature selection. We removed features with low variance, features that were blocklisted, features with missing values, and features with extreme outliers. Blocklisted features were those known to have caused issues in previous experiments and extreme outlier features were those that had a value greater than 15 standard deviations from the mean (Way, 2020). This procedure resulted in so-called “level 4” profiles. To form “level 5” consensus signatures, we collapsed level 4 replicate profiles into a single profile representing a compound-dose signature. For complete processing details, refer to https://github.com/broadinstitute/lincs-cell-painting. We further processed the Cell Painting data by performing 0-1 normalization on Cell Painting because not all features were on the same scale and we did not want different scales to influence the predictive power of a certain feature.

For L1000, we use the previously processed 978 “landmark” genes as our input features. We do not include all inferred genes because this would likely overload our VAE with redundant information. See (Subramanian *et al*, 2017) for complete processing details.

As input into our machine learning models, we split the data into an 80% training, 10% validation, and 10% test set, stratified by plate for Cell Painting and stratified by cell line for L1000.

### 5.3 Variational autoencoder implementations

A standard Vanilla VAE (Kingma & Welling, 2013) minimizes the loss for the sum of two loss functions: reconstruction (by mean squared error (MSE)) and Kullback–Leibler (KL) divergence. KL divergence encourages the latent space samples to follow a multivariate Gaussian distribution.

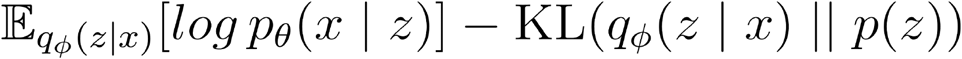

In a β-VAE, we multiply the KL divergence term by a constant β (Higgins *et al*, 2016). The purpose of this is to achieve greater disentanglement in the latent code. Our method of determining the value of β is described in the next section “5.4 Training procedure”.

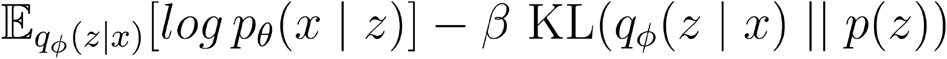

However, β-VAEs still suffer from two main problems. First is the “information preference property”, where if the KL divergence term is too high, all *z* values will be close to the prior p(x), leading to little useful information encoded in the latent space. The second problem is that if the regularization term is not strong enough, we get so-called “entangled” latent representations. These tradeoffs in a β-VAE led to the development of the maximum mean discrepancy (MMD)-VAE, where the scientists completely replaced the KL divergence term with a term that minimizes the mean maximum discrepancy (Zhao *et al*, 2019). Specifically, MMD forces the aggregated *z* distribution towards the prior rather than each individual *z* (as in the vanilla and β-VAE), which allows individual *z* values to diverge from the prior and more flexibly encode information (Wild, 2018). Also, just like how we use β to adjust the magnitude of the KL divergence term, we use λ to adjust the magnitude of the MMD regularization term for our MMD-VAE. We calculated MMD efficiently using the kernel embedding trick (Zhao).

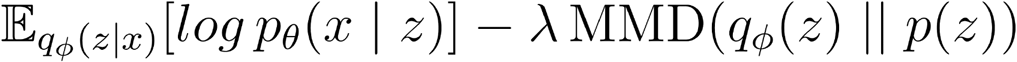

We used an Adam optimizer for all VAE architectures and datasets, and we used a leaky rectified linear activation function (Leaky ReLU) for all intermediate layers. Each model was a two-layer VAE, meaning that both encoder and decoder had one hidden layer. This hidden layer had 250 nodes for Cell Painting and 500 nodes for L1000 (determined by approximately one half of the number of input features).

### 5.4 Determining β in our β-VAE

The original β-VAE paper explains how to choose β. It states that if we have labelled data, we should use a disentanglement metric. But in our case where our data is not labelled, “the optimal value of β can be found through visual inspection of what effect the traversal of each single latent unit z has on the generated images (x|z) in pixel space.” But this also didn’t apply to our situation since we were not using images.

So, we proposed a new method of determining β by measuring the similarity between the original training set and simulated data points by sampling from a uniform distribution as is traditional in VAE generative models. To simulate data points that are close to the original distribution of the data, the VAE needs to have a balance of reconstruction and disentanglement. It needs good reconstruction because we want simulated data to look like real data. But we also need disentanglement because we simulate data points by sampling from the prior multivariate Gaussian, so the VAE benefits from the KL divergence term that pushes *z* values closer to that distribution.

It is likely that this method is not perfect and has a bias towards a lower-than-optimal β. This is because while simulating data benefits from a disentangled latent space, as long as the latent code roughly aligns with the prior, the simulation will be decent. On the other hand, performing LSA also requires that each feature be independent of each other; we need to be able to independently adjust the features to generate new unseen profiles. Because of this, the β determined by this method is likely to be lower than what would be optimal to perform LSA. Nonetheless, using this method would still give us a lot better β than if we were to choose randomly. Additionally, in practice, we observed that different β had a relatively little effect on cross validation performance.

Specifically, in our β-optimizing method, we measured similarity by calculating the Hausdorff distance between the original training data and the same number of simulated data points (Birsan & Tiba, 2006). Because simulating data points requires a balance of reconstruction and disentanglement—which is controlled by β—if we were to train the model many times with different values of β, we would observe that an intermediate β would result in the lowest Hausdorff distance. For L1000, β=40 was optimal. We found that for our Cell Painting datasets (level 4 replicate profiles and level 5 consensus signatures), a β < 1 was optimal (0.3 for Cell Painting level 5 and 0.06 for level 4). So, in this way, our implementation of β-VAE for Cell Painting differs from the original concept of β-VAE where you would increase β > 1to achieve disentanglement. For all three datasets, the magnitude of the KL divergence term (after multiplied by β) was between 10-30% of the total loss (MSE + KLD) (CP5 0.79/2.37 = 33%, CP4: 0.12/0.76 = 15%, L1000: 133/1330 = 10%).

We first chose a reasonable set of hyperparameters to use based on our initial training observations (latent_dim = 50, learning_rate = 0.001, encoder_batch_norm = True, batch_size = 128, epochs = 50) to keep constant and then adjusted β across many training sessions. After determining the optimal β (lowest Hausdorff distance), we used this value as a constant during a hyperparameter sweep to determine the optimal value for the other hyperparameters.

We used the same hyperparameters to train both our Vanilla VAE and MMD-VAE. This was because the data would remain the same between different VAE variants, it is likely that the same hyperparameters would perform well as well. Also, because hyperparameters can have a big impact on training performance, keeping them consistent would be the best way to compare these models. Furthermore, we did not observe much difference in cross validation performance for many combinations of hyperparameters we tested. When training the MMD-VAE, we first tried using the Hausdorff distance method to find the optimal value for λ. But, increasing λ didn’t have much effect on simulated data even as we increased λ much greater than1. So, we decided to choose λ merely by choosing a large value that still resulted in a stable training curve. This was 1000 for Cell Painting level 5, 10,000 for Cell Painting level 4, and 10,000,000 for L1000. For all three datasets, the magnitude of the MMD term (after multiplied by λ) was between 74-98% of the total loss (MSE + MMD) (CP5: 11.07/14.99 = 74%, CP4: 14.39/14.75 = 98%, L1000: 1948/2514 = 77%). This proportion of the regularization term is a lot higher than it was in the β-VAE. This ability for us to increase the magnitude of the regularization in a MMD-VAE with little negative consequences is a property of MMD-VAEs because their loss function allows them to encode information in their latent space even when the regularization term is high.

### 5.5 Hyperparameter optimization

Using Keras Tuner, we performed Bayesian hyperparameter optimization on all three datasets to select optimal learning rate (1e-2, 1e-3, 1e-4, 1e-5), batch size (32 to 512, with increments of 32), latent dimension (5 to 150, with increments of 5), and encoder batch normalization (True, False) for a two-layer VAE **(Table 1, Supplementary Figure 10)**. The optimal latent dimension of 10 for Cell Painting level 5 data was surprising as it implies that 10 features are sufficient to encode Cell Painting profiles.

**Table 1.** Hyperparameter combination of the top performing models for each dataset.

| Dataset | latent_dim | learning_rate | encoder_batch_norm | batch_size | val_loss |
| --- | --- | --- | --- | --- | --- |
| Cell Painting level 5 | 10 | 0.001 | True | 96 | 2.24 |
| Cell Painting level 4 | 90 | 0.0001 | True | 32 | 0.74 |
| L1000 | 65 | 0.001 | True | 512 | 1363.45 |

### 5.5 Cell Painting latent space feature interpretation

The optimal Cell Painting level 5 VAE models had 10 latent space features. Keeping all other latent space features at 0, we manipulated one feature at a time, comparing the reconstruction of +3 standard deviations with -3 standard deviations. Since σ of p(x) is 1, we used the values 3 and -3 for each latent space feature. To compare reconstructions, we took the absolute value of the difference between the two reconstructions. This output represents which original Cell Painting features most contribute to reconstruction of the single latent space feature. We repeated this procedure with all 10 latent space features.

### 5.6 Latent space arithmetic (LSA) approach to predict polypharmacology cell states

We transformed all 50,303 level 4 and 10,368 level 5 consensus profiles for the Cell Painting dataset, and 118,050 L1000 profiles into the latent space and then grouped them by their MOA annotation. We used the Drug Repurposing Hub MOA annotations (Corsello *et al*, 2017). To perform LSA, we first needed to filter compounds to include only those compatible with our hypothesis. Specifically, we included only compounds that satisfied the following rules. We only kept compounds with one or two MOA annotations. Also, for groups with only one annotated MOA, we only kept it if it corresponded to at least one group of compounds annotated with two MOAs. That is, if there was a group with MOA “A”, we only kept that group if there was another group with MOA “A|B” or “B|A”. The “|” notation indicates that the compound has evidence for both mechanisms. We also kept the DMSO profiles (negative control), which lacks MOA annotations. We needed to keep this group because it was part of our LSA equation hypothesis (“A” - “D” + “B” = “A|B”).

For each group, we calculated the mean for each latent space feature, giving us a vector of length 10 for each MOA combination. We included all doses of all annotated compounds to compute the mean latent space features. Then, for each existing “A|B” in our data, we performed vector addition and subtraction on the groups of MOA “A”, “B”, and “D” using our LSA equation, allowing us to have a predicted latent space representation of “A|B”. We then decoded this prediction to a reconstructed representation of the “A|B” and compared this representation with the original, real-data representation. The original representation for an MOA was achieved by taking the mean value of profiles with that MOA for all features. This comparison was done by computing the L2 distance for each MOA combination (see results).

To determine if the LSA approach was significantly different from a randomly shuffled control, we randomly shuffled all MOA labels, including DMSO labels, and performed the same LSA experiment. This results in a second distribution of L2 distances between the mean profile of an actual MOA label and a predicted profile from performing LSA on random profiles. We performed this shuffling 10 times to get a representative distribution of random predictions, so our control distribution was 10 times larger than the unshuffled distribution.

To compare against negative control baselines, we performed this same LSA procedure using principal component analysis (PCA) with 10 principal components. We also performed LSA using the original data dimensions, without any dimensionality reduction.

To determine the MOAs our VAE could predict the best, we calculated a z-test for each MOA combination between the 1) L2 distance between the predicted profile values and the actual mean profile values and 2) the distribution of L2 distances between the 10 predicted profile values from the control and the actual mean profile values. By doing this, we evaluated for polypharmacology MOA, A|B, how much better LSA with unshuffled VAE features could predict the profile values compared to the randomly permuted control distribution. A low test statistic would mean the LSA performed well for that particular polypharmacology prediction. A high test statistic indicates that the specific MOA combination could not be predicted, either because of incorrect annotations, non-additive or synergistic treatment effects, or a low penetrant phenotype.

### 5.7 Computational reproducibility

All scripts and computational environments to download and process data, train all VAEs, and reproduce all results in this paper can be found at https://github.com/broadinstitute/cell-painting-vae (Chow & Way, 2021).

## 6. Acknowledgements

We would like to thank Rachel Gesserman and Michael Mavros for their support and coordination of the Broad Summer Scholars Program (BSSP). This study was funded by the National Institutes of Health (R35 GM122547 to AEC).

